# MPC is a facultative pyruvate / lactate exchanger

**DOI:** 10.1101/2025.11.29.691055

**Authors:** Natali Cárcamo-Lemus, Daniela Rauseo, Alejandro San Martín, Carlos F. Lagos, L. Felipe Barros, Pamela Y. Sandoval

## Abstract

The mitochondrial uptake of pyruvate is mediated by the MPC. In the present study, we investigated the functional properties of the carrier using genetically encoded fluorescent sensors. Mitochondrial lactate efflux was inhibited by >80% by both pharmacological and genetic MPC ablation. MPC-dependent pyruvate influx accelerated the efflux of lactate, a fingerprint of shared transport. The non-metabolized pyruvate analog oxamate, which was found to enter mitochondria solely through the MPC, was also able to trans-accelerate lactate. Reportedly, mitochondria can produce lactate under reductive stress; our findings indicate that the MPC participates in this energy/redox venting by exchanging pyruvate by lactate.

## Introduction

Pyruvate is a major mitochondrial substrate, used for oxidative phosphorylation (OxPhos), cell growth, proliferation, and repair. The uptake of pyruvate by the organelle is chiefly mediated by the mitochondrial pyruvate carrier (MPC), plus an unclear contribution by monocarboxylate transporters (MCTs) (1,2). The MPC is structurally unrelated to the MCTs, has a higher affinity for pyruvate (1), and is not deemed to transport lactate (3,4). However, we recently observed that mitochondrial lactate uptake is partially sensitive to the MPC inhibitor UK5099 and to genetic ablation of the MPC. In addition, pyruvate was found to trans-accelerate lactate, although the relative contributions of the MPC and the MCTs were not addressed (5). Here we present unambiguous evidence that the MPC is also a lactate transporter.

## Results

### Mitochondrial pyruvate/lactate exchange

The MPC was approached in a native environment, as its kinetic properties are sensitive to partner proteins like ALDH4A1 (6). Substrate selectivity was first tested in HEK293 cells, a human cell line equipped with a robust OxPhos (7). Direct access to mitochondria was obtained by digitonin permeabilization of the plasma membrane under conditions that preserve respiration (5,8). Mitochondrial lactate and pyruvate levels were monitored with genetically encoded fluorescent sensors (**Figure 1A**). Lactate uptake was found to be insensitive to pharmacological and genetic MPC inhibition and failed to trans-accelerate matrix pyruvate, pointing to no MPC participation in the influx of lactate (**F1B-D**). Strikingly different results were obtained when the MPC was probed in the opposite direction. The efflux of lactate was sensitive to both strategies of MPC inhibition (**Figure 1E**), and pyruvate was effective at inducing lactate efflux, with an apparent K_M_ of 8 ± 4 µM, consistent with the high affinity of the MPC for pyruvate (9). No trans-acceleration was detected in the MPC knockout (**Figure 1F**). Trans-acceleration, also termed accelerated exchange, is considered the gold standard test for two substrates sharing the same transporter (10).

**Figure 1.**
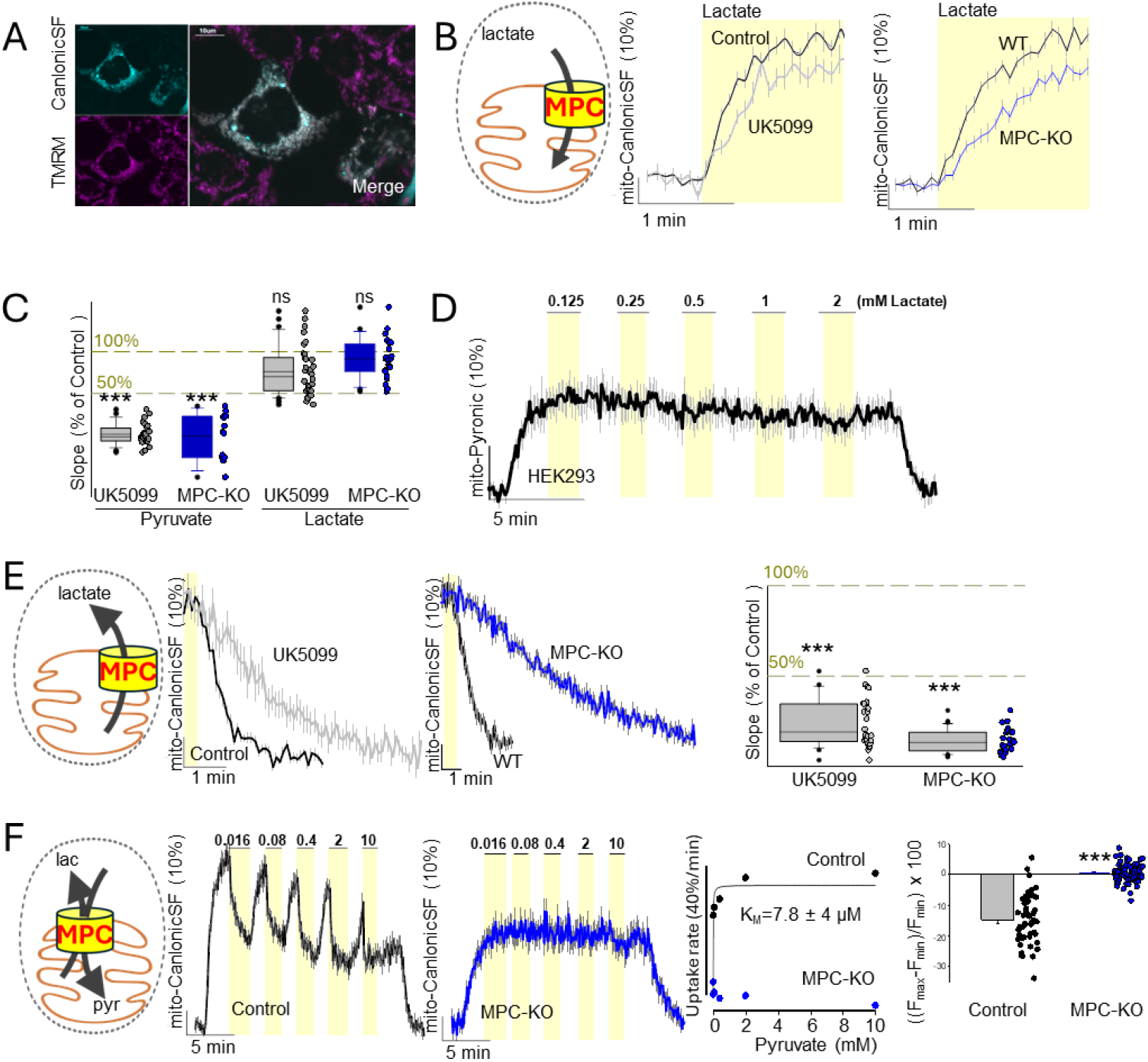
Mitochondrial pyruvate/lactate exchange. HEK293 cells were permeabilized with digitonin. **(A**) Colocalization of mito-CanlonicSF with TMRM-stained mitochondria. Size bar 10µm. (**B**) Lactate uptake in cells with pharmacological (UK5099) or genetic (MPC-KO) disruption of MPC. Representative data as the mean ± SEM (n = 10 cells). (**C**) Effect of MPC inhibition or deletion on initial pyruvate and lactate uptake rates. Data from three independent experiments, including 28 (UK5099) and 13 (MPC-KO) cells for pyruvate, and 34 (UK5099) and 24 (MPC-KO) cells for lactate. Each dot represents an individual cell. Mann-Whitney test; ***p < 0.001. (**D**) Trans-acceleration of mitochondrial pyruvate upon sequential lactate additions. (**E**) Mitochondrial lactate efflux under pharmacological or genetic MPC disruption. Data shown as mean ± SEM from n = 10 cells per condition. Normalized efflux quantification (ΔF·min^−1^). Data from three independent experiments from n = 26 and 30 cells. Unpaired t-test: ***p < 0.001. (**F**) Mitochondrial lactate efflux induced by pyruvate. Trans-acceleration rates were fitted to a hyperbolic decay model to estimate the apparent K_M_. Maximal fluorescence change at 80 µM. Data represent mean ± SEM of three independent experiments, and each dot corresponds to an individual cell. Unpaired t-test: ***p < 0.001.

### MPC is responsible for the pyruvate/lactate exchange

MPC inhibition almost eliminated pyruvate accumulation in the matrix (**Figure 1C**), but this result does not rule out MCT participation. Because pyruvate is metabolized in the matrix, its level is also sensitive to the energetic status of the organelle (**Figure 2A**). Thus, failure to detect pyruvate accumulation does not necessarily indicate that pyruvate influx is absent. To circumvent this limitation, we used oxamate, a non-metabolized pyruvate analog transported by the MPC and MCTs (11). Fortunately, we found that Pyronic also senses oxamate (**Figure 2B**) and that its uptake is completely abolished by genetic ablation or by pharmacological inhibition of MPC (**Figure 2C-D**). The absence of oxamate uptake when MPC function is lost indicates that mitochondria lack a significant second route of pyruvate import, supporting the use of oxamate as a specific probe for MPC activity. Figure **2E-F** shows that oxamate was capable of robustly trans-accelerating lactate efflux and that the trans-acceleration was abrogated by MPC disruption. Comparable results were obtained in an additional cell line (**Figure 2F**). Together, these findings demonstrate that the MPC is a lactate exporter. The MPC has recently been crystallized with the substrate-binding site either facing the intermembrane space (IMS) or the matrix. Docking simulations show that lactate can bind to both conformations (**Figure 2G-H**). Strikingly, the affinity score for lactate was even better than that for the known MPC-substrate oxamate.

**Figure 2.**
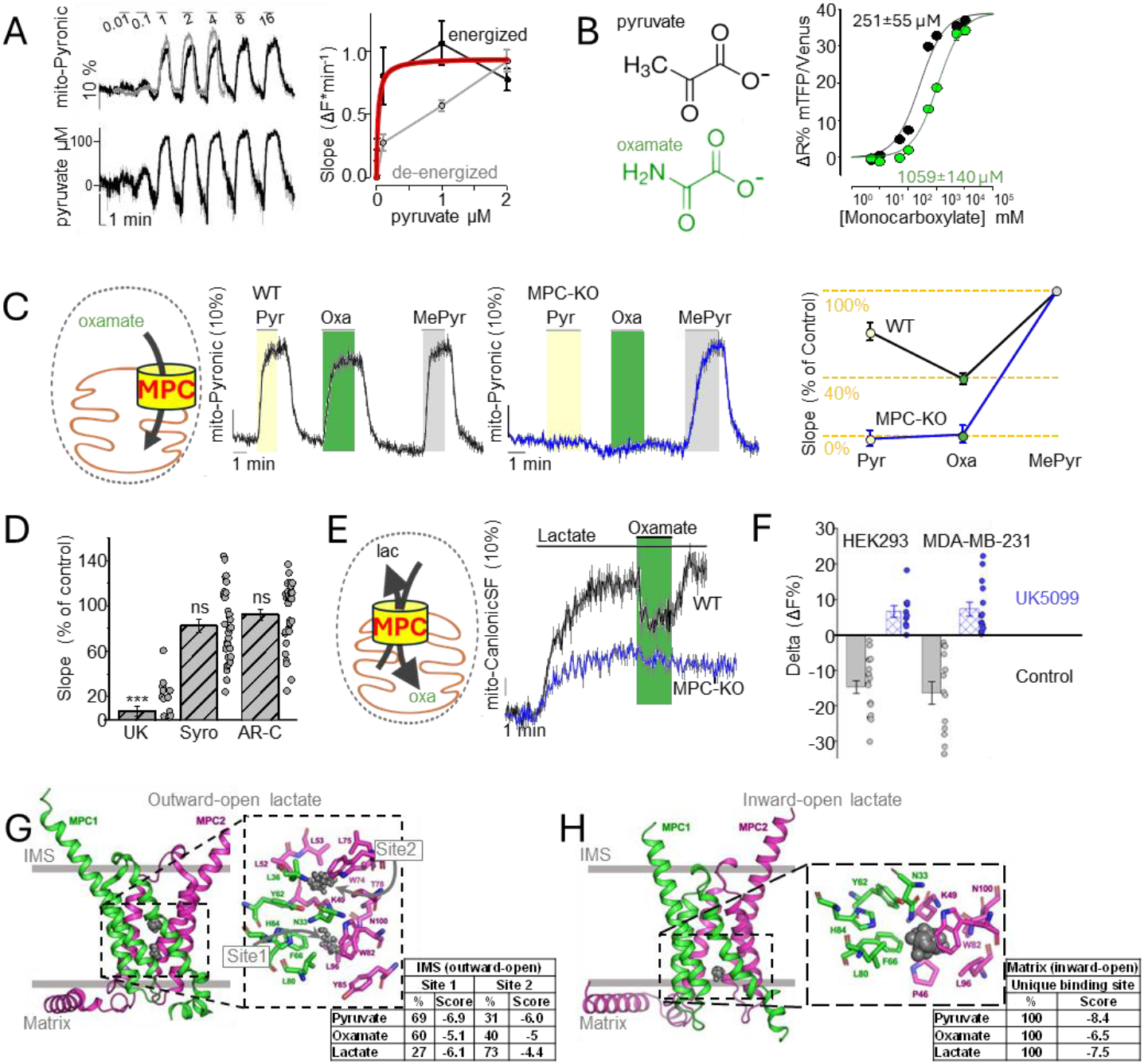
Validation of lactate exchange with a non-metabolizable MPC substrate. (**A)** Permeabilized cells under high redox (black) or de-energized (gray) conditions were exposed to increasing pyruvate concentrations. Fluorescence changes and initial uptake rates were fitted to a hyperbolic model, yielding apparent K_M_ values of 44 ± 30 µM (energized, n = 2, 14 cells) and 970 ± 560 µM (de-energized, n = 3, 12 cells). Data are mean ± SEM. (**B**) In vitro kinetics of purified Pyronic response to increasing pyruvate or oxamate concentrations. Apparent K_M_ values were calculated in triplicate. Curves were fitted to a one-site saturation model; data shown as mean ± SEM. (**C**) WT and MPC-KO cells expressing mito-Pyronic were sequentially exposed to pyruvate (Pyr), oxamate (Oxa), and methyl-pyruvate (MePyr). Uptake rates were normalized to methyl-pyruvate (control) and expressed as mean ± SEM (n = 3). (**D**) Mitochondrial pyruvate uptake rates in cells treated with 1 µM UK5099 (UK, 21), 5 µM syrosingopine (Syro, 30), or 1 µM AR-C155858 (AR-C, 30), normalized to untreated control. Data from three experiments; each dot represents one cell. Statistics: UK5099 (t-test, p < 0.001), syrosingopine and AR-C155858 (Mann–Whitney, not significative). (**E**) Trans-acceleration of mitochondrial lactate with 6 mM oxamate in HEK293 wild type (WT) and MPC-deficient (MPC-KO) cells. Data presented as mean ± SEM (10 cells, representative of three experiments). (**F**) Trans-acceleration of lactate by 6 mM oxamate in mammalian cells. ΔF expressed as % of the lactate steady state (mean ± SEM, n = 3). (**G**) Monocarboxylate docking outward-open MPC. Lactate centers of mass from 100 simulations are represented by grey dots, flanked by the two subunits of the MPC. Coordinating residues are indicated. Scores showed that lactate was found either in a high-affinity site (27 out of 100) or in a low-affinity site (73 out of 100). The table summarizes simulations for pyruvate, oxamate, and lactate. (**H**) Monocarboxylate docking inward-open MPC. Lactate centers of mass from 100 simulations are represented by grey dots, flanked by the two subunits of the MPC. Coordinating residues are indicated. Scores showed that lactate was found in a single high-affinity site (100 out of 100). The table summarizes simulations for pyruvate, oxamate, and lactate.

## Discussion

The main claim of this study is that the MPC operates as a facultative pyruvate/lactate exchanger. This conclusion is based on the observed pyruvate/lactate and oxamate/lactate exchange, and on the dramatic effect of pharmacological and genetic MPC inhibition on both lactate efflux and exchange.

The notion that the MPC does not transport lactate (12) originated from experiments with purified mitochondria, in which cold pyruvate trans-accelerated the efflux of radiolabeled pyruvate, but lactate was without effect (11,13). Soon after, a thorough kinetic analysis by one of the groups reported trans-acceleration of pyruvate efflux, but only at supraphysiological lactate (14). These results and the discovery of UK5099 as a blocker of mitochondrial pyruvate uptake that does not target MCTs led to the notion that the MPC is a different protein (11,15). Eventually, the cloning of the MPC revealed that this carrier belongs to a different gene family (3,4). We could not find any recent study addressing the lactate selectivity of the MPC.

The crystal 3D structure of the MPC was reported in 2025 (16–19). Lactate is not mentioned in any of the four studies. The key element for substrate and UK5099 recognition is the carboxylic group, in register with efficient transport of phenylpyruvate, acetoacetate, and dichloroacetate (11,13,14). This property also helps to explain why lactate is also a substrate. According to our docking simulations, lactate binds better at the matrix-open conformation of the carrier, which might help to explain why the MPC specializes in lactate efflux. A non-exclusive explanation for the functional asymmetry is that lactate can enter mitochondria via a parallel pathway (5), likely dissipating the driving force for pyruvate extrusion. In contrast, pyruvate relies solely on the MPC as entry pathway.

What is the function of lactate extrusion by the MPC? Our group has recently described a phenomenon termed mitochondrial energy venting (5), in which the exchange ability of the MPC appears instrumental. When there is little lactate about, the MPC works as a standard co-transporter, driven by the trans-mitochondrial proton gradient and pyruvate removal by oxidative metabolism. Under reductive conditions like hypoxia, matrix NADH rises, and now pyruvate is converted into lactate, which is exchanged by cytosolic pyruvate via the MPC. The pathway is futile in terms of carbons but loses one reducing equivalent per transport cycle, helping to alleviate reductive stress and diminish ROS production (5). Lactate also drives the lactylation of multiple matrix proteins, including LDH and other metabolic enzymes (5), with unknown functional consequences.

The MPC is a transporter of relevance for metabolic disorders and chronic diseases like diabetes and cancer. Another contribution of this work, with translational potential, is the realization that oxamate is sensed by Pyronic. A non-metabolized probe is useful to avoid the confounding effects of metabolism in high-throughput screening assays for drug discovery.

## Material and Methods

### Reagents

Original constructs used in this work were: Pyronic (Addgene #51308), CanlonicSF (gifted by Alejandro San Martín; Addgene plasmid # 178342), and pLentiCRISPR-v1-sgMPC2_7 (gifted by Jason Cantor; Addgene plasmid # 163457).

### Cell cultures

MDA-MB-231, HeLa, and HepG2 cells were obtained from the American Type Culture Collection (ATCC). MDA-MB-231 cells were cultured at 37°C in L-15 (Lebovitz) media without CO_2,_ whereas HEK293, HeLa, and HepG2 cells were maintained at 37°C in a humid atmosphere with 5% CO2. HEK293 cells were grown in DMEM/F-12 (1:1), HeLa cells in high-glucose DMEM, and HepG2 in DMEM low-glucose. All media were supplemented with 10% fetal bovine serum (FBS). For superfusion assays, cells were seeded onto poly-L-lysine-coated glass coverslips and studied at 40-60% confluence.

### Production of the HEK293 cell lines stably expressing a metabolic sensor

Genetically encoded fluorescent indicators Pyronic and CanlonicSF were engineered to include (4X) mitochondrial targeting sequence derived from subunit VIII of human cytochrome c oxidase. In the case of mito-CanlonicSF, the sensor was expressed bicistronically with a mitochondria-targeted mCherry, providing an internal fluorescence reference for normalization of mitochondrial signal intensity. The mito-Pyronic, mito-CanlonicSF, and Laconic were independently packaged into lentiviral particles and transduced into HEK293 cell lines at a multiplicity of infection (MOI) of 1. Blasticidin (10 µg/mL) was used for the selection of stably transduced cells, and individual clonal lines were subsequently isolated by fluorescence-activated cell sorting (FACS) using a BD FACS Jazz Cell Sorter.

### MPC2-KO generation

HEK293 cells stably expressing mito-Pyronic or mito-CanlonicSF were cultured at 40% confluency in a 6-well plate. Lipofectamine 3000 (ThermoFisher #L3000015) was used for the transfection of pLentiCRISPR-v1-sgMPC2_7 to disrupt MPC2 via CRISPR-Cas9. After 48 hours, puromycin was added at 10 µg/ml for the selection of transfected cells. Puromycin-resistant clones were expanded and cultured for four or five passages and then used in perfusion assays.

### Baculoviral particle production for the transient expression of metabolic sensors

Recombinant baculoviral particles encoding the mitochondrial or cytosolic targeted sensor constructs were produced in-house using the Bac-to-Bac Baculovirus Expression System (Thermo Fisher Sci). Briefly, Sf9 cells were transfected with purified bacmid/sensor DNA using CellFECTINII reagent to generate the initial viral stock (P1). Viral amplification was performed through successive infections in Sf9 cells to obtain high-titer P3 stocks. Viral supernatants were harvested 72 h post-infection, clarified by centrifugation (500 × g, 10 min), filtered through a 0.45 µm membrane, and stored at 4°C until use. The baculoviral dose yielding ∼95% transduction efficiency in HEK293-FT cells was empirically determined and used for subsequent infections.

### Imagen acquisition

An upright Olympus FV1000-confocal microscope equipped with solid-state laser lines and a 20× water-immersion objective (N.A.1.0) was used to collect time series images every 3 or 5 seconds in XYT scan mode (scan speed: 156 Hz; 800 × 800-pixel; pinhole 800 μm). Mito-Pyronic, Pyronic, and Laconic were excited at 440 nm, and emissions were collected simultaneously at 480 ± 15 nm and 550 ± 15 nm. Mito-CanlonicSF was excited at 488 nm with emission collected at 525 ± 25 nm. A bicistronically expressed red fluorescent protein was used for normalization of mito-CanlonicSF and excited at 543 nm and detected at 610 ± 50 nm. The fluorescent signal for a region of interest (ROI) from each cell was collected. Background subtraction was performed with ImageJ.

### Mitochondrial Staining

Mitochondria were labeled with TMRM (tetramethylrhodamine methyl ester perchlorate; Thermo Fisher Scientific), a cationic fluorescent dye that selectively accumulates in mitochondria of live cells. Cells were incubated with 40 nM TMRM in KRH buffer (in mM: 136 NaCl, 3 KCl, 1.25 CaCl_2_, 1.25 MgSO_4_, 10 HEPES, pH 7.4) at room temperature directly under the microscope objective. Time-lapse images were collected every few seconds until optimal visualization of the mitochondrial network was achieved.

### Kinetic assays

Intact cells were constantly superfused at room temperature with KRH buffer. Permeabilization of cells was previously described in Rauseo et al. (2025). In brief, cultures were perfused for 3 min with 30 µM digitonin and 200 µM ADP in intracellular buffer (in mM: 130 KCl, 10 NaCl, 1.25 MgCl_2_, 0.37 CaCl_2_, 1 EGTA, 10 HEPES, pH 7.2, and 284-286 mOsm/L). For digitonin removal, 10 min of perfusion with intracellular buffer was applied before recordings. All permeabilized kinetic assays were performed under energized conditions unless otherwise indicated. Mitochondrial energization was achieved by supplementing the intracellular buffer with 0.2 mM glutamate and 0.1 mM malate. Monocarboxylate transport was assessed using 10 mM pyruvate or 1 mM lactate as substrates. For MPC inhibition, UK5099 was applied at 1 µM. For substrate comparison assays, pyruvate, oxamate, and methyl-pyruvate were each applied at a final concentration of 5 mM.

### MPC trans-acceleration assays

Mitochondria from permeabilized cells were preloaded with 0.5 mM pyruvate and trans-accelerated with 0.125, 0.25, 0.5, 1, and 2 mM lactate. Alternatively, mitochondria preloaded with 0.5 mM lactate were then trans-accelerated with different concentrations of pyruvate (0.016, 0.08, 0.4, 2, and 10 mM). The trans-acceleration of mitochondrial lactate was also assessed using 6 mM oxamate.

### Docking-based binding analysis

The cryo-electron microscopy structures of the apo human MPC in the matrix-facing state (PDB id 8YW8) at a resolution of 3,1Å was used as the starting point (18). Nanobodies and small molecule ligand cardiolipin were deleted. The MPC1/2 structure was prepared using ChimeraX (UCSF) to add hydrogen atoms and assign the amberff99sb charges (20). A receptor grid of 15×15×15 Å was prepared using FRED receptor. The ligands were constructed using Marvin Sketch v5.7.0 (ChemAxon Ltd. Hungary) and saved as multimol2 file. Multiconformer library of compound libraries was prepared using OMEGA v6.1.0.1 (OpenEye Scientific Software, Santa Fe, NM). QUACPAC v1.6.3.1 (OpenEye Scientific Software) was used to assign the AM1BCC charges to each compound conformer. Molecular docking of the compounds was performed using FRED v4.3.3.1 (OpenEye Scientific Software, Santa Fe, NM) (21). Candidate poses of the ligands within the receptor site (100) were obtained and optimized using the Chemgauss4 scoring functions. Consensus structures of the poses returned from exhaustive docking, and optimization will be obtained by consensus scoring using the PLP, Chemscore and Chemgauss3 scoring functions (22). Predicted binding affinity was obtained using the LUDI empirical scoring functions (23). In the matrix-open state, lactate forms hydrogen bonds with N33 and Y62 of MPC1 and K49 and N100 of MPC2, with its hydroxyl group engaging Y62. Pyruvate overlays with essentially the same geometry and interaction network, while oxamate occupies the same site, preserves the core contacts, and introduces an additional interaction with H84 of MPC1.

### Statistical analysis

Data analysis was performed in SigmaPlot-12 software (Jandel). A paired t-test was applied to the before-and-after protocols with normal distribution, p < 0.001. Alternatively, an unpaired t-test was used to compare independent groups, ***p < 0.001. Normality was assessed using the Shapiro-Wilk test; data failing this criterion were analyzed with the Mann-Whitney-Wilcoxon signed rank test (pairs) ***p < 0.001.

## Acknowledgments

We thank the members of the Energy Metabolism Group at CECs for helpful discussions. This work was funded partly by FONDECYT projects 1230145 (LFB), USS-FIN-23-FAPE-03 (PYS), and FONDEQUIP EQY240004 (LFB).

## Notes

### Competing Interest Statement

The authors have declared no competing interest.

